# Integrative Identification and Characterization of PCOS-Associated lncRNAs From the Interface of Genetic Association, Transcriptomics, and Gene Structure Evolution

**DOI:** 10.64898/2026.03.31.715548

**Authors:** Zhizhou He, Yibai Li, Tatiana P. Shkurat, Elena V. Butenko, Ekaterina G. Derevyanchuk, Svetlana V. Lomteva, Li Chen, Leonard Lipovich

## Abstract

**Background:** Polycystic ovary syndrome (PCOS) is a prevalent endocrine disorder and a leading cause of female infertility, with complex genetic, metabolic, and hormonal etiologies. Long non-coding RNAs (lncRNAs) have emerged as important regulators of diverse biological processes, yet their roles in PCOS remain underexplored. Here, we identified and characterized PCOS differentially expressed gene-associated lncRNAs (PDEGAL) with an integrative approach combining expression data, genetic association, and evolutionary analysis.

**Methods:** Thirty-three PCOS-associated protein-coding genes were obtained from our prior study, and all their nearby and overlapping lncRNAs were annotated. These candidates were analyzed using UCSC Genome Browser-mapped annotations and datasets, including NCBI RefSeq, GENCODE, GTEx, GWAS SNPs, and conservation, as well as the FANTOM5 cap analysis of gene expression (CAGE) promoter data, to assess their expression, regulatory potential, genetic variant overlaps, and evolutionary conservation.

**Results:** Twenty-three PDEGALs (18 antisense to, and 5 sharing bidirectional promoters with, known PCOS-associated protein-coding genes) were identified. 17 PDEGALs contained GWAS SNPs with statistically significant disease associations, 9 of which were associated with PCOS-related traits. 5 PDEGALs demonstrated expression in the KGN granulosa cell model of PCOS. Key gene structure element (KGSE) analysis revealed that most PDEGALs are primate-specific. Integrating four criteria—GTEx expression, GWAS SNPs, FANTOM promoterome, and KGSE conservation—highlighted HELLPAR as the only lncRNA fulfilling all four, while five others—PGR-AS1, MTOR-AS1, ENSG00000265179, ENSG00000256218, and LOC105377276—fulfilled three of the four criteria.

**Conclusions:** We have systematically identified candidate PCOS regulatory lncRNAs with convergent genetic, expression, and evolutionary evidence. These results provide a framework for functional validation and highlight lncRNAs as potential biomarkers and therapeutic targets in PCOS that function by regulating their nearby and overlapping protein-coding genes.

## 1. Introduction

Polycystic ovary syndrome (PCOS) is a complex endocrine disorder affecting up to 20% of women of reproductive age worldwide, and it accounts for approximately 80% of female infertility cases [1,2]. The PCOS diagnosis was first described by Stein and Leventhal in 1935, and the condition is currently diagnosed using the Rotterdam criteria, requiring the presence of at least two of the following: hyperandrogenism, oligo-or anovulation, and polycystic ovarian morphology [3,4]. The pathogenesis of PCOS is interconnected with multiple conditions, including chronic low-grade inflammation, insulin resistance, genetic susceptibility, and lifestyle factors. Notably, 40–85% of PCOS patients are overweight or obese, and insulin resistance is present in up to 70% of obese and 30% of overweight PCOS patients [5,6]. In addition to reproductive dysfunction, PCOS significantly increases the risk of pregnancy-related disorders, including gestational diabetes, preeclampsia, and preterm birth [7-9]. The central features of PCOS include follicular dysregulation and hyperandrogenism, which is linked to granulosa cell (GC) dysfunction, including apoptosis and aberrant gene expression [10,11]. Several therapeutic modalities for PCOS management have become available, such as ovulation induction, inositol supplementation, and assisted reproductive technologies. However, they focus on managing symptoms rather than curing the disease. Consequently, long-term or lifelong therapeutic management remains the standard approach for treating PCOS [12,13]. This highlights the need to study PCOS at the molecular level, especially considering the relevance of recently identified high-level disease regulatory factors, including long non-coding RNAs (lncRNAs), to its etiology. The resulting insights could lead to new ways to treat the disease.

The ENCODE (Encyclopedia of DNA Elements) Consortium, the direct successor of the Human Genome Project, revealed that the human genome consists of a vast amount of non-coding sequences, while only approximately 2% of the genome encodes proteins [14,15]. Growing evidence indicates that the human genome is transcribed into a vast repertoire of noncoding RNAs, most of which are not functionally annotated yet. In the FANTOM (Functional Annotation of Mammalian cDNA) Consortium, we demonstrated that the mouse genome has more noncoding-RNA genes than protein-coding genes, a finding later confirmed in the human genome [14,16-18]. Among the non-coding transcripts, long non-coding RNAs (lncRNAs)—defined as RNA molecules longer than 200 nucleotides lacking open reading frames—are the most abundant class and have emerged as newly recognized critical regulators of gene expression at both transcriptional and post-transcriptional levels [14,19]. LncRNAs can be categorized into biotypes based on their genomic location and orientation relative to protein-coding genes (including antisense RNAs, long intergenic non-coding RNAs, intronic transcripts), as well as by functional characteristics (including enhancer RNAs, telomere-associated RNAs, circular RNAs)[14,20-23]. We were the first group to describe the genome-scale lack of conservation of lncRNAs at and near protein-coding gene loci between human and mouse, the prevalence of “coding-noncoding” (coding-lncRNA) gene pairs of both the sense-antisense and the bidirectional-promoter gene pair classes globally, and the frequent incidence of “gene chains” uniting 3 or more genes within a locus through multiple sense-antisense overlaps and bidirectional promoters [18]. We subsequently demonstrated that cis-antisense lncRNA transcription exerts locus-specific, both positive and negative, regulatory effects on the protein-coding (sense) genes occupying the same loci, suggesting multiple possible regulatory mechanisms [24] that can also be relevant in endocrine and reproductive pathologies including PCOS.

LncRNAs contribute to a broad range of essential biological processes, including chromatin remodeling, X-chromosome inactivation, transcriptional regulation, genomic imprinting, and nuclear transport. Dysregulation or mutation of lncRNAs has been implicated in the development and progression of numerous human diseases, including cancers, cardiovascular disorders, and endocrine abnormalities [25-27]. In recent years, accumulating evidence, including from our work on coordinated differential expression of sense-antisense, coding-noncoding gene-lncRNA pairs in the human myometrium during pregnancy [28], has highlighted the broad relevance of lncRNAs to reproductive health and disease [28-30]. Moreover, PCOS studies in mouse and pig models pinpointed abnormal lncRNA expression profiles in the serum, ovaries, adipose tissue, and follicular fluid [31-33]. Despite these advances, only a limited number of PCOS-related lncRNAs have been functionally characterized, highlighting the need to further investigate lncRNAs’ specific roles in ovarian dysfunction and reproductive outcomes.

In a pioneering analysis in this field, Diao et al. (2004) constructed a cDNA library and, using custom microarray hybridization, identified 290 transcripts that were differentially expressed in ovarian cells from PCOS patients relative to controls, including only 119 from known protein-coding genes [34]. Subsequent studies demonstrated that long non-coding RNAs (lncRNAs) include numerous high-level network regulators that directly control the expression of microRNAs and protein-coding genes. LncRNAs can regulate the expression of their nearby or overlapping protein-coding genes through several cis-regulatory mechanisms; this observation provides a rationale for investigating gene-overlapping and gene-proximal lncRNAs in the context of PCOS - in particular, lncRNAs that reside at or near protein-coding genes that are differentially expressed and/or functionally implicated in PCOS pathogenesis. While lncRNA genes comprise the majority of human genes, most human lncRNA genes are primate-specific and – in contrast to protein-coding genes - are not conserved in other mammals [27,35,36]. To date, no study has integratively examined PCOS-associated lncRNAs by combining genome-wide association studies (GWAS)-driven discovery of candidate genes, disease-relevant tissue-specific transcriptome analysis, and evolutionary conservation. Our study aims to fill this gap by implementing a comprehensive approach to identify and characterize lncRNAs that serve as putative proximal regulators of PCOS-associated protein-coding genes.

## 2. Results

## 2.1. Twenty-three lncRNAs were identified at the genomic loci of thirty-three protein-coding? genes with known PCOS functions

We initiated our search using 33 protein-coding genes established as key drivers of PCOS pathogenesis in human granulosa cells [37]. These 33 foundation genes were curated based on their direct involvement in ovarian granulosa cell functions, specifically focusing on hormone receptor signaling (FSHR, LHCGR, and AMHR2), the PI3K-AKT-mTOR and PI3K-AKT-Foxo3a intracellular pathways—which are essential for follicle activation—and steroidogenesis, particularly aromatase synthesis. Through systematic analysis of their genomic associations with lncRNAs, we found that 14 of them overlap antisense lncRNA genes, three shared bidirectional promoters with immediately-adjacent lncRNA genes, and four exhibited both features (Figure 1, Table 1 and Table S1). In the latter category, the lncRNA PACERR is antisense to the PTGS2 gene transcript isoform ENST00000680451.1_2 and shares a bidirectional promoter with the PTGS2 transcript isoform NM_000963.4; we classified it as an antisense lncRNA. There were 23 PDEGAL lncRNAs (Table 2). Eight of these 23 lncRNAs have NCBI RefSeq gene model support, indicating robust, unambiguous expression and gene structure evidence. These NCBI-curated RefSeq lncRNA genes are: PDE4B-AS1 (NR_123718.1), MTOR-AS1 (NR_046600.1), LIF-AS1 (NR_149070.1), LIF-AS2 (NR_148946.1), FGF10-AS1 (NR_108034.1), LOC105377276 (NR_188415.1), PACERR (NR_125801.1), and LOC101929319 (NR_110248.1). Notably, we identified complex “gene-chain” architectures in four loci [18]. Specifically, the MTOR, PDE4B, and TNFAIP6 gene chains combine the protein-coding gene with two lncRNA genes and exhibit a dual configuration featuring bidirectional promoters (BDP) as well sense-antisense (S-AS) overlaps, whereas LIF displays a distinct ‘2x S-AS’ structure involving two different antisense transcriptional units overlapping the same protein-coding gene (Figure 2A-D).

**Table 1.** Known PCOS-related protein-coding genes and lncRNAs at and near their loci.

| PCOS related genes | Antisense lncRNAs | Bidirectional promoter sharing lncRNAs |
| --- | --- | --- |
| 33 | 18 | 5 |

**Table 2.** PDEGAL lncRNAs and their PCOS-related partner protein-coding genes. The first 18 lncRNAs are antisense to PCOS related genes, and the last 5 lncRNAs share bidirectional promoters with PCOS related genes. Strandedness is per the hg19 (GRch37) human genome assembly, as viewed in the UCSC Genome Browser (accessed between January and June 2025).

| LncRNA | LncRNA strand | Coding gene | Coding strand |
| --- | --- | --- | --- |
| ENST00000607600.1_2 | - | VEGFA | + |
| PGR-AS1 | + | PGR | - |
| PDE4B-AS1 | - | PDE4B | + |
| ENSG00000268650 | + | PDE4C | - |
| ENSG00000258430 | + | AKT1 | - |
| MTOR-AS1 | + | MTOR | - |
| GCLC-AS1 | + | GCLC | - |
| ENSG00000265179 | - | ADCYAP1 | + |
| ENSG00000287848 | - | AKR1C4 | + |
| LIF-AS1 | + | LIF | - |
| LIF-AS2 | + | LIF | - |
| HELLPAR | + | IGF1 | - |
| FGF10-AS1 | + | FGF10 | - |
| ENST00000743643.1_1 | - | ALAS1 | + |
| LOC105377276 | - | EREG | + |
| PACERR | + | PTGS2 | - |
| LOC101929319 | - | TNFAIP6 | + |
| ENST00000520598.1_2 | + | STAR | - |
| ENST00000760442.1_1 | - | PDE4B | + |
| ENST00000838067.1_1 | + | MTOR | - |
| ENSG00000284633 | + | PDGFB | - |
| ENSG00000256218 | - | NTF3 | + |
| ENSG00000289267 | - | EGR1 | + |

**Figure 1.**
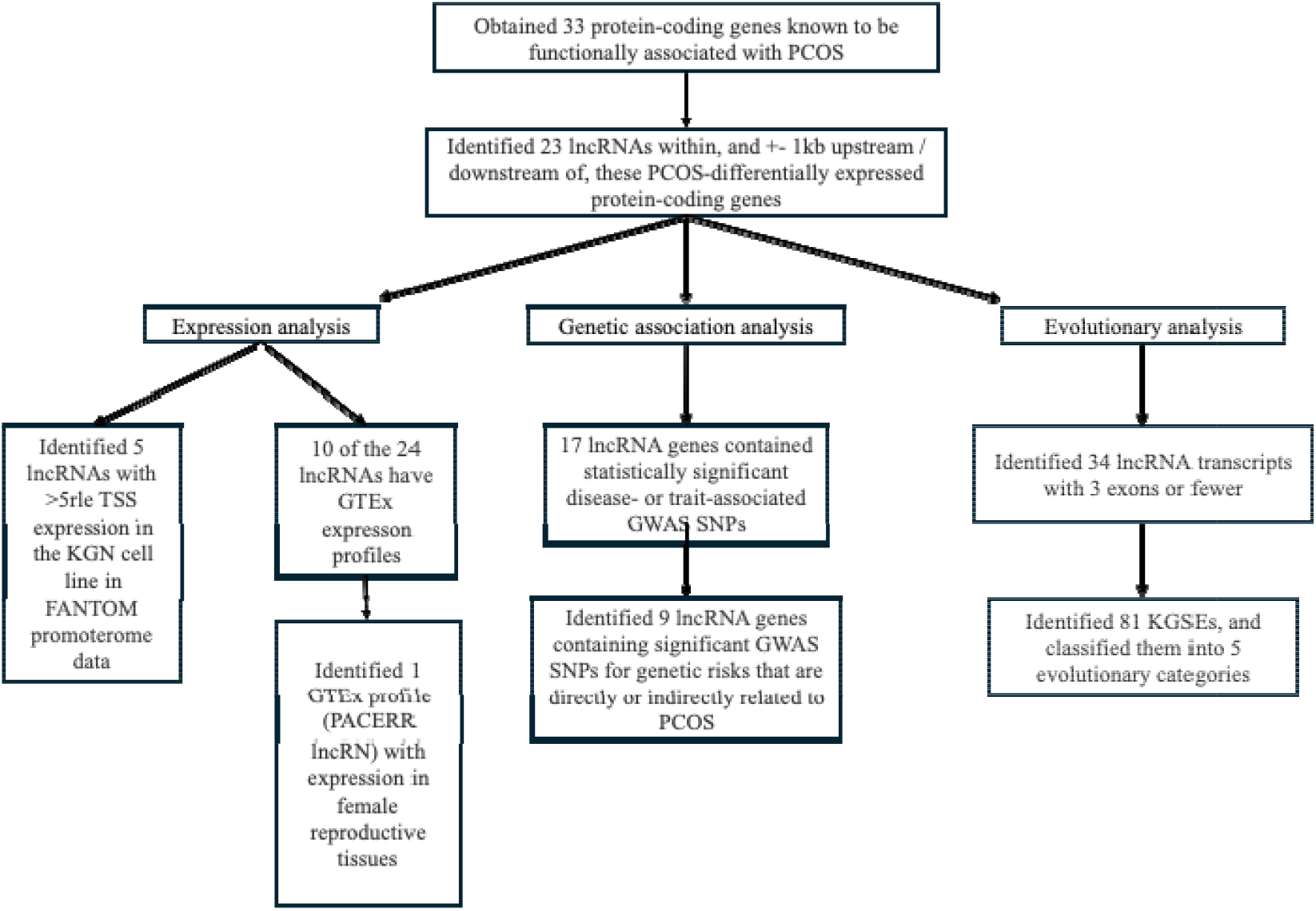
Flowchart of PDEGAL characterization.

**Figure 2.**
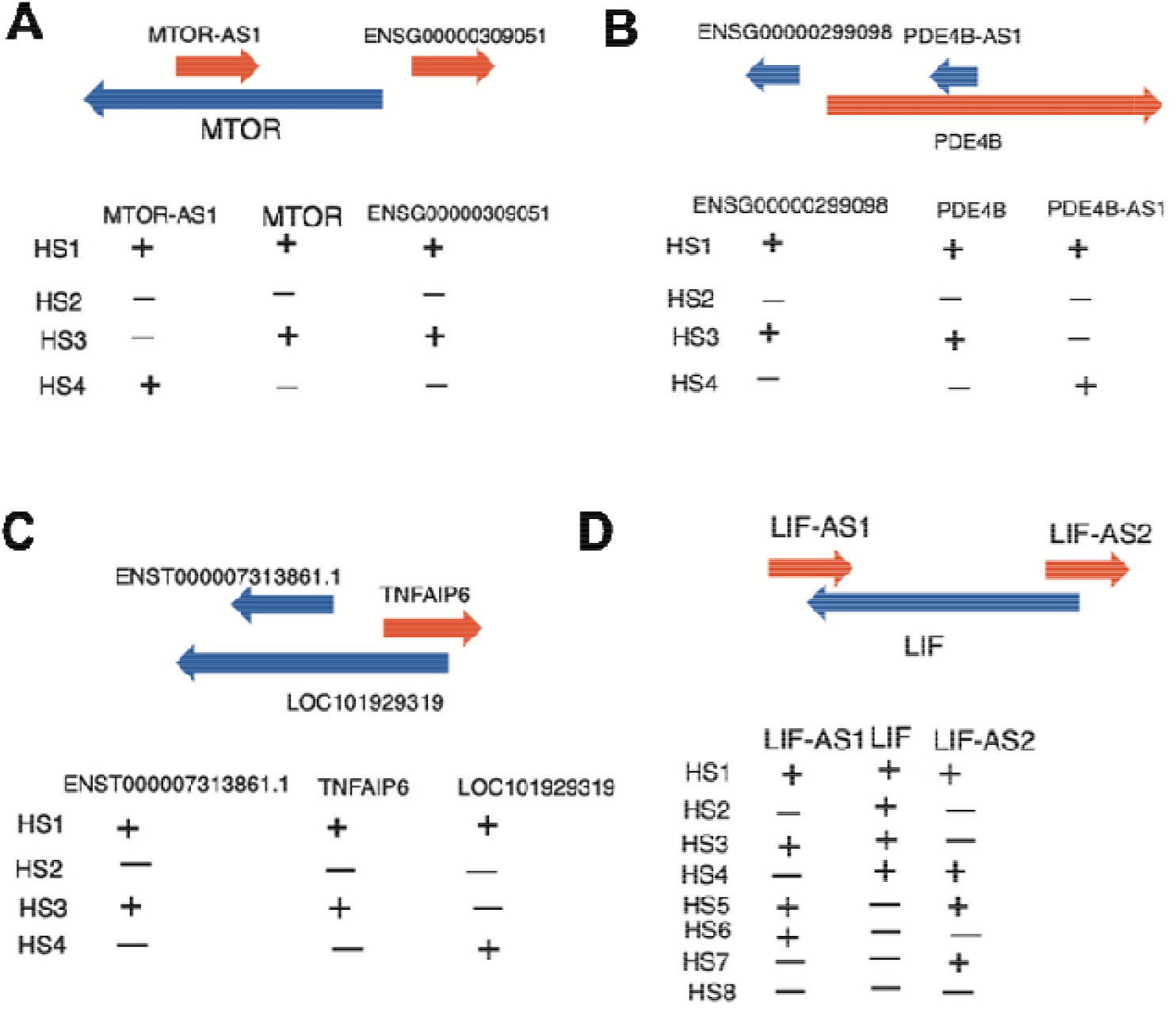
Complex “gene-chain” architectures at PCOS-associated genomic loci. (A–C) The MTOR, PDE4B, and TNFAIP6 loci exhibit a dual configuration, characterized by the presence of both a shared bidirectional promoter (BDP) and sense-antisense (S-AS) transcript overlap. (D) The LIF locus displays a distinct ‘2x S-AS’ architecture, involving multiple discontinuous regions of antisense overlap. HS: Hypothetical State (of gene chain expression). The symbols (+) and (−) indicate the possible expression states of the transcripts. Genes joined by bidirectional promoters are assumed to be coregulated. Genes forming sense-antisense pairs are assumed to be capable of either positive or negative regulation. Gene structures and orientations are based on UCSC Genome Browser visualizations.

### 2.2. The lncRNA PACERR is expressed in female reproductive tissues

We analyzed 33 PCOS-related protein-coding genes and identified 23 PDEGAL lncRNAs, which were further classified according to the type of evidence: 10 lncRNAs had GTEx data, 9 contained PCOS-related GWAS SNPs, 5 were expressed in KGN cells in the FANTOM promoterome, and 22 contained key gene structure elements (KGSEs) (Table S2). HELLPAR was the only lncRNA satisfying all four criteria. Additionally, five lncRNAs—PGR-AS1, MTOR-AS1, ENSG00000265179, ENSG00000256218, and LOC105377276—fulfilled three of the four criteria. From the 10 GTEx-represented lncRNAs, they were well-supported by RNAseq in this reference collection of human transcriptomes from over 50 tissues of 948 individuals. However, only PACERR demonstrated an expression level of > 1 TPM (transcripts per million) in reproductive tissues. PACERR is detectable in GTEx ovary, uterus, cervix, and vagina transcriptomes, and it is also expressed in other tissues, including the prostate and colon (Figure 3). In contrast, several other PDEGAL lncRNAs - PDE4B-AS1(NR_123718.1), GCLC-AS1(ENST00000655377.1_3), ENSG00000265179(ENST00000581719.3_3), and ENSG00000265179 (ENST00000582554 1 4) – are only detected in testis in GTEx data While this might indicate their irrelevance to PCOS, GTEx-listed genes are biased in favor of RefSeq-annotated, highly expressed genes, and GTEx negative results do not exclude the possibility of PCOS-relevant expression at low levels or in tissues and cell types not profiled by that project.

**Figure 3.**
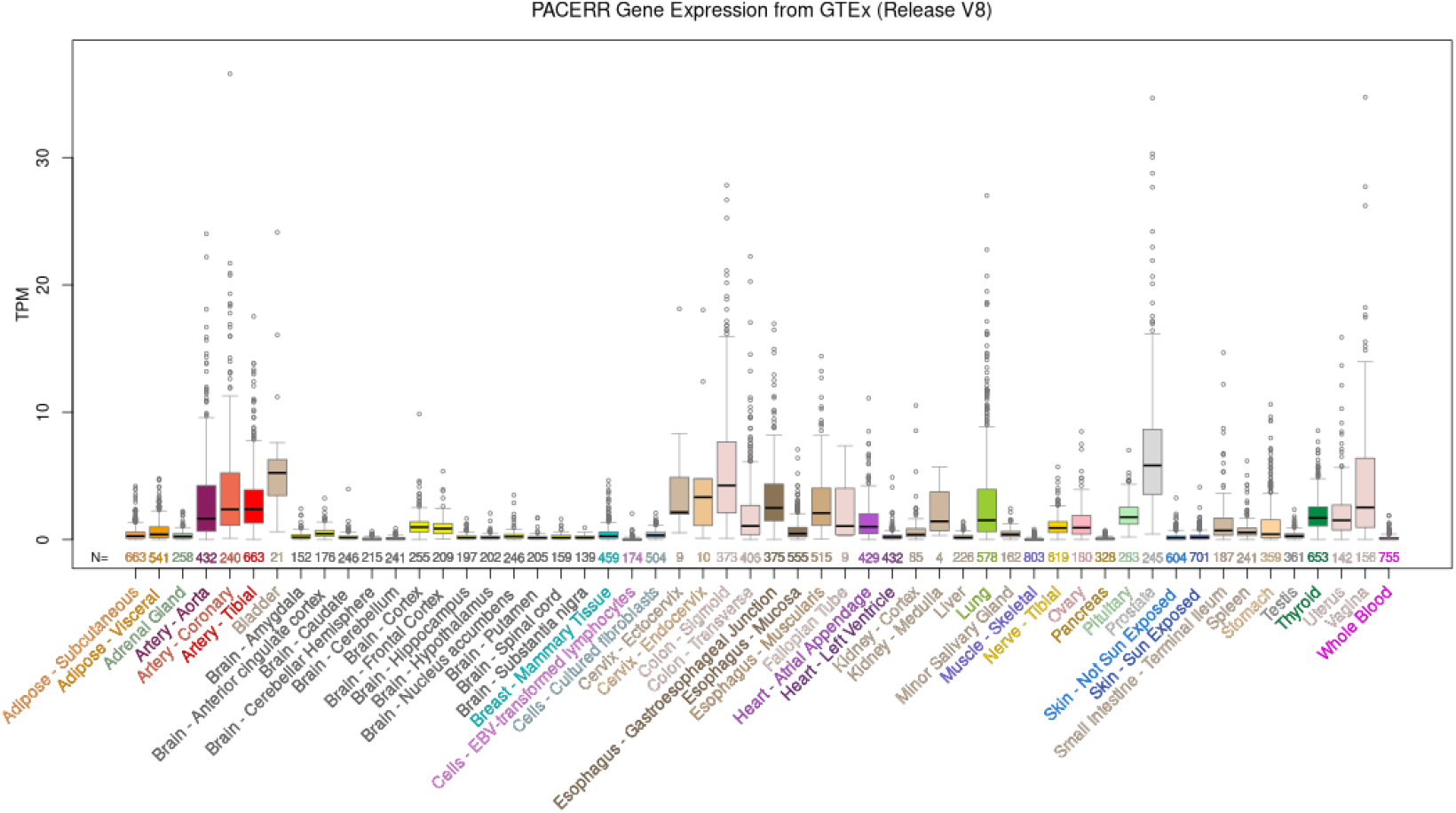
Screenshot of PACERR lncRNA expression in GTEx-profiled tissues. This lncRNA is expressed in several key female reproductive organs and tissues, as well as elsewhere.

### 2.3. Ten PDEGAL lncRNAs contain GWAS SNPs related to PCOS

Of the 23 PDEGAL lncRNA genes, 17 contain significant disease-or trait-associated GWAS SNPs from the NHGRI-EBI GWAS Catalog track of the UCSC Genome Browser. 9 of the 17 lncRNA genes harbored GWAS SNPs showing genetic risk association with female reproductive disorders, and with common diseases and quantitative traits that increase PCOS risk or are frequent PCOS co-morbidities, including obesity, body mass index, type 2 diabetes, insulin resistance, and hormonal traits (Table 3). These lncRNAs were PGR-AS1, ENSG00000268650, MTOR-AS1 (NR_046600.1), ENSG00000265179, ENSG00000287848, ENSG00000256218, HELLPAR (ENST00000626826.1_2), LOC105377276 (NR_188415.1), and LOC101929319 (NR_110248.1). To clarify the potential for direct functionality of these SNPs, we analyzed their specific locations within the lncRNA gene structures (Table 3), showing that not all of these SNPs were located in introns. Only LOC101929319 harbored an intronic SNP. In contrast, four lncRNA genes—PGR-AS1, MTOR-AS1, ENSG00000256218, and LOC105377276— contained exonic SNPs, which are more likely to represent variants that are directly functional rather than merely serving as genetic markers of risk conveyed by other sequences within their linkage disequilibrium (LD) blocks. Furthermore, SNPs in five lncRNAs—ENSG00000268650, MTOR-AS1, ENSG00000265179, ENSG00000287848, and LOC101929319—were located within the sense-antense overlap region between the lncRNA gene and its protein-coding partner. These significant, NHGRI- and EBI-catalogued GWAS SNPs are suggestive of PCOS-associated genetic risks mapping to the PDEGAL lncRNAs they reside within. Although the localization of all these SNPs within the PDEGAL lncRNAs shows the lncRNAs to be the closest genes to the SNPs (distance of zero), and therefore makes the lncRNAs direct causal candidates for functionally contributing to the conditions the SNPs are significantly associated with, it is also possible that the SNPs’ disease associations are due to the nearby PCOS-associated protein-coding genes or other genes in their LD blocks.

**Table 3.**
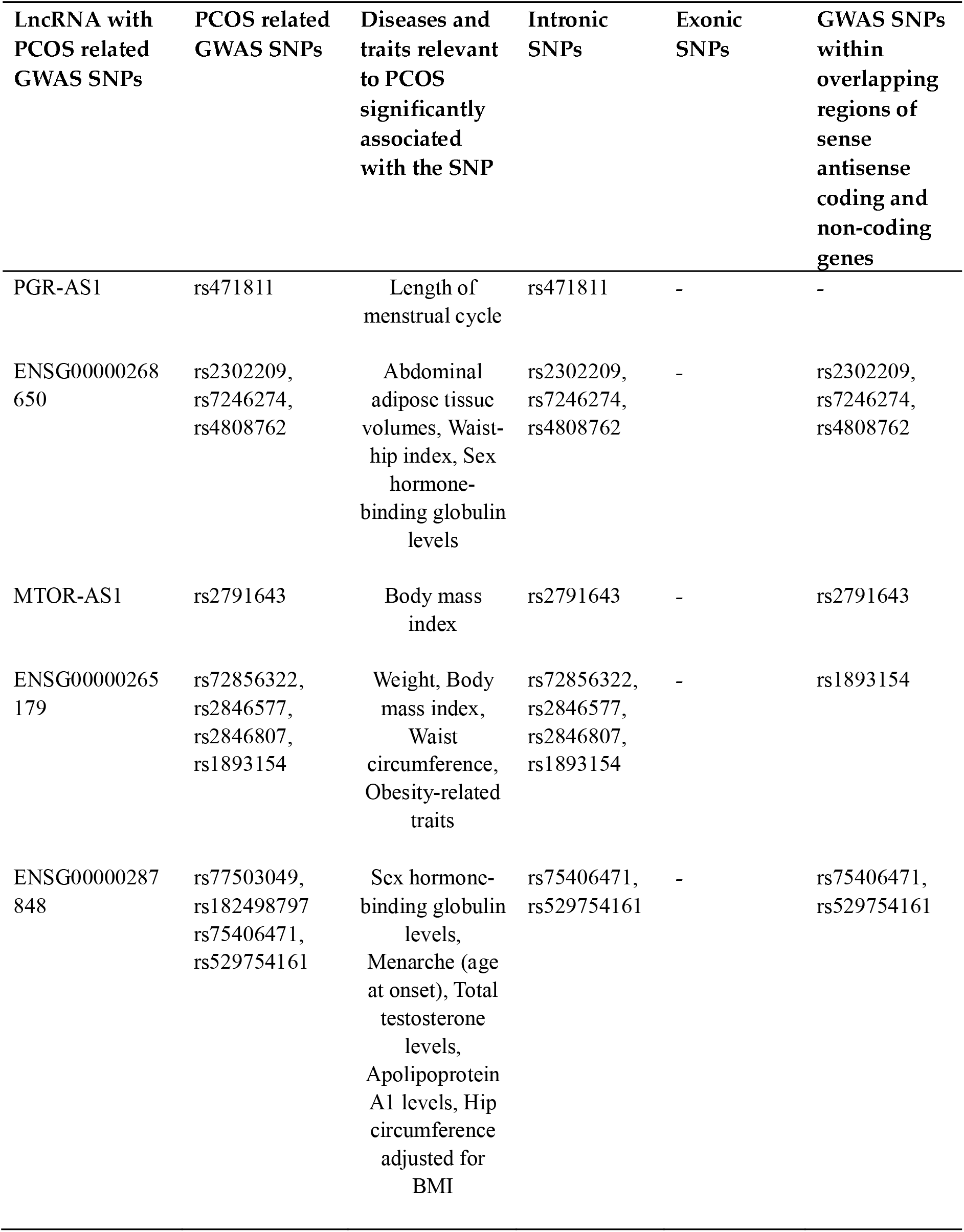
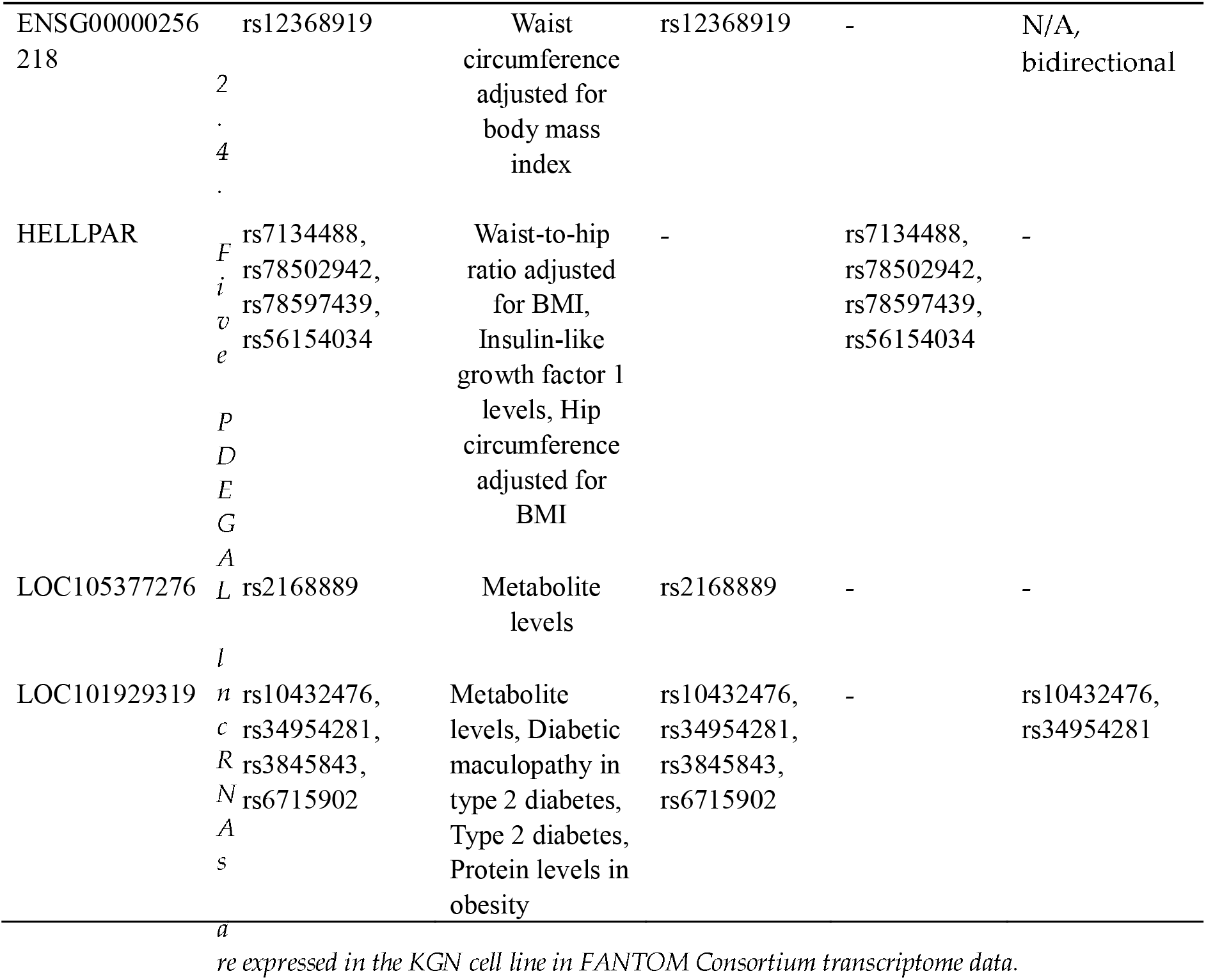
PDEGAL lncRNA genes containing GWAS SNPs significantly associated with traits or conditions directly related to PCOS. SNP identifiers, and the diseases and traits the SNPs are significantly associated with, are displayed. The table also details whether these SNPs are intronic, exonic, or in sense-antisense overlap regions.

Using the FANTOM5 hg19 human promoterome view of the ZENBU Genome Browser [38], we analyzed the genomic regions containing the 23 PDEGAL lncRNAs, focusing on the TSS (transcription start site) of each lncRNA. We identified 5 lncRNAs expressed in the ovarian granulosa cell tumor cell line KGN, a commonly used in-vitro PCOS model, in the conservative CTSS and less-restrictive BAM FANTOM CAGE data (Table 4). These lncRNAs represent feasible targets for functional validations in the widely available KGN cell line. Their knockdown or overexpression should allow interrogation of their regulatory relationships with their overlapping or nearby PCOS-associated protein-coding genes in future work.

**Table 4.** LncRNAs with >5rle KGN expression in CAGE Phase1 CTSS and CAGE Phase1 BAM.

| lncRNA with KGN expression | KGN level in CAGE Phase1 CTSS (rle) | KGN level in CAGE Phase1 BAM (rle) |
| --- | --- | --- |
| PGR-AS1 | 0.429 | 6.446 |
| ENSG00000284633 | 0 | 5.157 |
| ENSG00000256218 | 0 | 5.587 |
| HELLPAR | 0 | 5.157 |
| LOC105377276 | 0 | 56.229 |

### 2.5. KGSE evolutionary analysis of PDEGAL lncRNAs pinpoints recent and primate-specific gene structure changes

To assess the evolutionary origin timing of Key Gene Structure Elements (KGSEs), defined as the GT-splice donors, -AG splice acceptors, and AATAAA or ATTAAA polyadenylation signals, of these lncRNA genes, and to determine whether these sequence features may have undergone recent changes after the mammalian radiation or after the primate-nonprimate divergence, we performed KGSE evolutionary conservation analysis of PDEGAL lncRNA genes. The Gould and Brosius gene-birth model stipulates that de novo originations of promoters and KGSEs over evolutionary time may drive gene birth [39]. Examining KGSEs in multispecies genome alignments allows the relative dating of each KGSE’s origin, and its evolutionary changes, relative to lineage and species common ancestors, determining the probable timing of the origin, and of any major gene-structure changes, of the gene containing the KGSEs under analysis [40]. We examined the conservation of each key gene structure element (KGSE) of PDEGALs by searching for conservation of the splice sites at the start and the end of each intron (GT and AG respectively) and of the polyadenylation signal consensus sequence near the 3’end of the last exon (AATAAA or ATTAAA). First, we noted the closest (to human, along the species phylogeny) non-human species in which the KGSE KGSE is not conserved (“near” means that the species was a primate, “far” means that it was a nonprimate). Then, we noted the most distant (from human) non-human species in which the KGSE was conserved (“near” means that the species was a primate, “far” means that it was a nonprimate).

We focused on 34 transcripts belonging to 12 PDEGAL lncRNA genes that met our predefined criterion of containing three or fewer exons. We evaluated the interspecies conservation of 81 KGSEs from these 34 transcripts, comprised of 17 polyadenylation signals, 31 splice donors (19 Intron 1 GT and 12 Intron 2 GT), and 33 splice acceptors (21 Intron 1 AG and 12 Intron 2 AG), in the 99-species MultiZ alignment (hg19 assembly). 15 KGSEs were classified as near–near, 39 as near–far, 22 as far–far, 3 as N/A–near, and 2 as N/A–far, based on their conservation profiles across vertebrate species. The N/A category (N/A-near and N/A-far), which accounted for 5 of the 81 KGSEs, indicates that no nearest non-conserved species was found; in these cases, the KGSE is well conserved in all species in which it exists. A prevalent trend was that the “near-far” pattern was the most frequent, outnumbering other KGSEs in splice donors, splice acceptors, and polyA signals. Notably, all PolyA signals from single-exon PDEGAL transcripts were classified as near-far. This suggests recent sequence diversification of these KGSEs in primates, despite the ancient origin of the KGSEs. We also assessed the relative age of splice sites, finding that splice donors appear slightly younger than splice acceptors in this set of lncRNA genes: among 31 splice donors, 8 had a “near” furthest conserved species compared to 23 “far,” while among 33 splice acceptors, only 6 were “near” compared to 27 “far.” Finally, the nearest nonconserved species for all KGSE types were more likely to be primates, suggesting that the majority of these transcripts with pre-primate origins did not begin to structurally vary until recent primate lineages. Furthermore, a detailed co-evolutionary analysis of 35 introns based on their paired splice donor-acceptor categories (Table S3) revealed a heterogeneous distribution of patterns, with the most frequent types being Far far - Near far (N=7), Near far - Near far (N=6), and Near far - Far far (N=6). We provide a representative example of phylogenetic diagram–based KGSE conservation analysis for the lncRNA PDE4B-AS1 (Figure S1), illustrating the method used to determine the nearest non-conserved and furthest conserved species.

## 3. Discussion

Here, we identified, from public transcriptome data and gene catalogs, 23 candidate lncRNA genes that are antisense to, and/or share bidirectional promoters, with protein-coding genes known to be functionally relevant to PCOS. Among these lncRNAs, PACERR deserves prioritization as a PCOS causal candidate, because it exhibits antisense overlaps as well as bidirectional promoter sharing with distinct PTGS2 isoforms, while also being expressed in multiple female reproductive tissues in the GTEx transcriptome repository, supporting its potential involvement in reproductive pathophysiology.

To further understand the functional as well as the genetic basis for the possible association of the PDEGAL lncRNAs with PCOS, we incorporated GWAS data on all statistically significant disease- and trait-associated SNPs residing within these lncRNA genes. 17 of these 23 lncRNAs contained significant GWAS SNPs, and for 9 of these, at least some of the SNPs were associated specifically with PCOS-related phenotypes, including reproductive disorders, obesity, BMI, insulin resistance, and type 2 diabetes. These genetic associations support the potential relevance of these lncRNAs as candidate causal contributors to PCOS etiology, pointing to them as candidates for future mechanistic studies.

Of the 17 PDEGAL lncRNAs containing statistically significant GWAS SNPs, 9 of which harbored SNPs that are significantly associated with PCOS-related traits, only PACERR showed appreciable expression (≥1 TPM) in female reproductive tissues in the GTEx data. The lack of concordance between significant GWAS SNP localization and appropriate tissue-specific expression of the SNP-harboring gene is not unprecedented, and is indicative of a challenge in functional genomics: the presence of a disease-associated SNP in a transcript does not necessarily imply transcriptional activity in the disease-relevant tissue context, at least in primary tissues under normal physiological conditions and in cancer cell line models that tend to be profiled in public transcriptome resources.

While 8 PDEGALs contain PCOS-relevant GWAS SNPs but lack PCOS-relevant tissue expression profiles, 7 other PDEGALs are supported by GTEx but not by GWAS. Several factors may explain this discrepancy, including context-specific expression, such as disease-or cell state-dependent activation, post-transcriptional regulation, or epigenetic silencing of these transcripts in the healthy tissues profiled by GTEx. Alternatively, the lncRNAs may function through cis-acting mechanisms on their nearby PCOS-functionally-associated protein-coding genes through mechanisms that do not require high levels of transcript abundance.

To investigate the feasibility of exploring these lncRNAs in a commonly used in vitro PCOS cell line model, we analyzed the FANTOM5 promoterome expression data. Five of the 24 lncRNAs exhibited transcription start site (TSS) activity in the KGN ovarian granulosa cell tumor cell line, with a relative log expression (rle) of ≥ 5, suggestive of their active transcription in a granulosa cell context. This is consistent with the hypothesis that these lncRNAs are functionally relevant in the ovarian cumulus granulosa cell environment which gives rise to PCOS.

Complementing the population-genomics and transcriptome-mining approaches to relating PDEGALs to PCOS, we performed evolutionary analysis of KGSEs (GT-AG splice sites and polyadenylation signals) to better understand the evolutionary origins, and changes, of the PDEGALs through conservation and divergence patterns of their fundamental gene structures. From the 34 transcripts with ≤3 exons, corresponding to 12 PDEGALs, 81 KGSEs (31 splice donors, 33 splice acceptors, and 17 polyA signals) were analyzed for conservation in 100 vertebrate species using the UCSC Genome Browser Multiz alignment track (Table 5). The results reflect a spectrum of evolutionary ages among these elements. The prevailing KGSE category (15) was “near-far,” meaning that the nearest species to humans where the KGSE is not conserved is a primate, while the furthest species from humans where the KGSE is conserved is a nonprimate mammal. This indicates that, despite a relatively ancient origin (in the common ancestor of primates and nonprimate mammals), PDEGALs’ KGSEs frequently undergo evolutionary change within the primate lineage. This may be due to selective pressures that finetune the lncRNAs’ gene structures in specific primate lineages, and/or interspecies differences in primates’ reproductive systems. Of the 81 KGSEs analyzed, 63 were either near–far or far–far, suggesting that the majority of PDEGALs’ KGSEs originated before the primate-nonprimate divergence (the second “far” means that the species furthest from human in terms of evolutionary distance, where the KGSE is conserved, is a nonprimate). 60 out of 81 KGSEs exhibited conservation within primate lineages, encompassing the near–near, near–far, and N/A–near categories. No credible nonmammalian conservation of any PDEGAL KGSEs was observed in the MultiZ alignments.

**Table 5.**
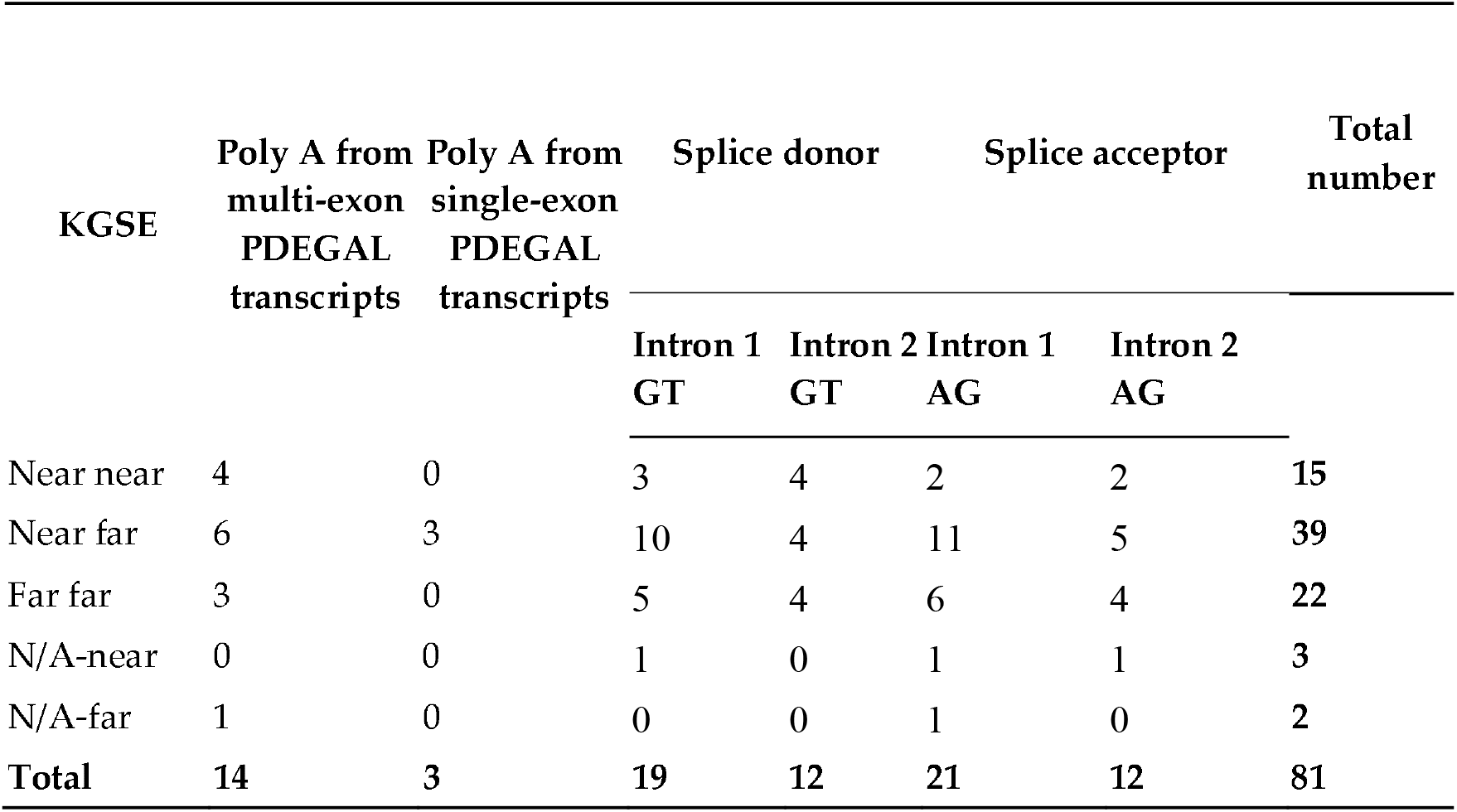
Evolutionary conservation analysis of 81 KGSEs in 34 PDEGAL lncRNA transcripts. Near-near suggests site conservation only in primates; Near-far indicates origin of the site before the primate-nonprimate split, with sequence diversification in primates and conservation in non-primate mammals; Far-far indicates deep conservation throughout, and possibly outside of, mammals. The N/A category (N/A-near, N/A-far) indicates that no nearest non-conserved species was found; in these cases, the KGSE is conserved in all species where it is detectable.

| KGSE | Poly A from<br>multi-exon<br>PDEGAL<br>transcripts | Poly A from<br>single-exon<br>PDEGAL<br>transcripts | Splice donor |  | Splice acceptor |  | Total<br>number |
| --- | --- | --- | --- | --- | --- | --- | --- |
|  |  |  | Intron 1 | Intron 2 | Intron 1 | Intron 2 |  |
|  |  |  | GT | GT | AG | AG |  |
| Near near | 4 | 0 | 3 | 4 | 2 | 2 | 15 |
| Near far | 6 | 3 | 10 | 4 | 11 | 5 | 39 |
| Far far | 3 | 0 | 5 | 4 | 6 | 4 | 22 |
| N/A-near | 0 | 0 | 1 | 0 | 1 | 1 | 3 |
| N/A-far | 1 | 0 | 0 | 0 | 1 | 0 | 2 |
| **Total** | **14** | **3** | **19** | **12** | **21** | **12** | **81** |

Our findings should facilitate experimental testing of these lncRNAs’ functions in granulosa cells and PCOS pathology, with methods including CRISPR genome editing, RNA interference, and reporter assays in the KGN cell line model as well as in primary patient-derived cumulus granulosa cells. These experiments should determine whether these lncRNAs may positively or negatively regulate the PCOS-associated protein-coding genes at their shared complex loci, and test whether the knockdown, overexpression, promoter modification, or deletion of the lncRNAs impacts PCOS-relevant cellular phenotypes. LncRNAs with PCOS-relevant tissue expression profiles that also contain PCOS-related GWAS SNPs should be prioritized for such *in vitro* functional studies. Our disease-candidate lncRNA annotation strategy also presents a model for systematically uncovering the contributions of gene-proximal and gene-overlapping non-coding RNAs to the etiology and molecular pathogenesis of other diseases.

## 4. Materials and Methods

### 4.1. Identification of PCOS-related genes

We obtained the 33 PCOS-associated protein-coding genes from our prior study [37]. We performed a bioinformatics screen using the UCSC Genome Browser (GRCh37/hg19 assembly) and identified all known protein-coding genes functionally associated with key granulosa cell processes involved in PCOS pathogenesis, including hormone receptor signaling (FSH, LH, AMH), activation of the PI3K-AKT-mTOR signaling pathway, and aromatase activity.

### 4.2. UCSC Genome Browser tracks

In the UCSC Genome Browser [41,42], we selected the human GRCh37/hg19 genome assembly. To visualize protein-coding genes and lncRNA transcripts, we used the GENCODE V48lift37 track under Genes and Gene Predictions in Pack mode [16], the GENCODE Versions track under Genes and Gene Predictions in Show mode [43], the NCBI RefSeq track under Genes and Gene Predictions at pack mode [44,45], and we hid all other tracks in Genes and Gene Predictions.

To visualize significantly disease-associated and trait-associated GWAS SNPs [46], we displayed the GWAS Catalog track, under Phenotypes, Variants, and Literature, in Full mode, and we hid all other tracks in Phenotypes, Variants, and Literature. After clicking on each GWAS SNP within the lncRNA sequence, we reviewed the displayed evidence (publications, annotations, specific statistically-significant disease, phenotype, and trait associations) to identify diseases and quantitative traits directly related to female reproductive disorders, obesity and body mass index, type 2 diabetes and insulin resistance, and hormonal traits including testosterone levels and sex hormone-binding globulin (SHBG). To visualize the expression of lncRNA in 54 human tissues of 948 individuals [47,48], we displayed the GTEx Gene V8 track, under Expression, in Pack mode.

### 4.3. FANTOM5 CAGE analysis of PDEGAL expression in KGN

The FANTOM Consortium mapped the human promoteromes of 1,000 primary cell and tissue types at a single-base resolution, by implementing the HeliScope Cap Analysis of Gene Expression (CAGE) approach [49,50]. CAGE Transcription Start Sites (CTSS) data points to the exact nucleotide where transcription begins, and also shows how frequently that TSS is used, measured in tags per million (TPM) in each tissue’s and cell type’s transcript library [49]. Binary Alignment Map (BAM), an alternate method of CAGE data interpretation, encompasses the raw alignments of CAGE reads to the genome and is less restrictive, allowing the detection of certain TSSs missed in CTSS. We examined the FANTOM5 CAGE Phase1 CTSS human tracks and CAGE Phase1 BAM human tracks for the presence of transcription start sites at PDEGALs in the KGN granulosa cell tumor cell line. We copied and pasted the lncRNA’s chromosomal location into the search bar of FANTOM (since both the UCSC Genome Browser and the FANTOM ZENBU Browser in our work used the hg19 assembly), and clicked “search”. We expanded the “Entrez gene hg19” track, “UCSC_gencodeV10_hg19_20120101” track, and “UCSC_RefSeq_hg19_20120101” track in the ZENBU Browser for visualization of the sense and antisense transcripts, and ensured that it matched the same transcripts’ display in the UCSC Genome Browser. Then, we expanded the “FANTOM5 CAGE Phase1 CTSS human tracks pooled filtered with 3 or more tags per library and rle normalized” track and scrolled down to locate the TSS expression in granulosa cell tumor cell line KGN, ensuring that the TSS signal matched the strand of the target lncRNA. We did the same for the “FANTOM5 CAGE Phase1 BAM human with Q3 filter and rle normalized” track.

### 4.4. Data collection, analysis, and interpretation

We canvassed the phylogenetic history of Key Gene Structure Elements (KGSEs), including all canonical splice site motifs (GT at the 5′ splice site and AG at the 3′ splice site of each intron) and polyadenylation signals (AATAAA or ATTAAA within the last 100 bp of the last exon), which together define the fundamental genetic structure of a gene. In 2013, we pioneered the implementation of this approach to examine evolutionary conservation of gene structures rather than gene sequences [40]. Here, we applied it to characterize the evolutionary conservation of gene structures of the lncRNAs that we identified at PCOS-associated protein-coding loci.

To investigate the evolutionary age and conservation of lncRNA transcripts, we used the 100-species Multi Z alignment to examine the presence and absence of KGSEs across species. The most phylogenetically distant species containing the KGSE was referred to as the “furthest conserved species,” representing the maximum known relative evolutionary age of that element. Conversely, the least phylogenetically distant species to humans that lacked the conserved KGSE was labeled the “nearest nonconserved species,” reflecting the evolutionary footprint of recent evolutionary changes that modified or abrogated that particular KGSE in a particular species or lineage.

We classified each KGSE into one of five categories, based on these two evolutionary indicators. We defined the indicator “near” as referring to nonhuman primate species and the indicator “far” as referring to nonprimate species in the UCSC Genome Brower’s 100-species MultiZ genome alignment. For each datapoint, the first evolutionary indicator describes the nearest nonconserved species, and the second evolutionary indicator describes the furthest conserved species. To keep the dataset size manageable, we focused on lncRNA transcripts with 3 or fewer exons. The biological meaning of the five categories can be summarized as follows:

- near–near: the first “near” means that the KGSE, though present in human, was recently lost (through sequence substitution abolishing the KGSE) in at least one nonhuman primate (nearest-to-human species where the KGSE is not conserved is “near,” i.e. a primate), and the second “near” means the KGSE is conserved only within primates and is hence of a recent primate-specific origin (furthest-from-human species where the KGSE is conserved is “near,” i.e. also a primate).
- near–far: the first “near” means that the KGSE, though present in human, was recently lost (through sequence substitution abolishing the KGSE) in at least one nonhuman primates (nearest-to-human species where the KGSE is not conserved is “near,” i.e. a primate), and the “far” means this KGSE originated before the primate-nonprimate divergence (furthest-from-human species where the KGSE is conserved is “far,” i.e. a nonprimate mammal).
- far–far: the first “far” means that the nearest-to-human species where the KGSE is not conserved is a nonprimate mammal (i.e. the KGSE is conserved in all primates, but exhibits change in at least one nonprimate), and the second “far” means that the KGSE originated early in mammalian evolution and is conserved widely across mammals. KGSEs in this class are completely conserved in all primates.
- N/A–near: the first “N/A” means that there is no “nearest non-conserved” species, because the KGSE was conserved across all alignable species. Therefore, there is only a “furthest conserved” species, which in this case, was a primate (“near”).
- N/A–far: the first “N/A” means that there is no “nearest non-conserved” species, because the KGSE was conserved across all alignable species. Therefore, there is only a “furthest conserved” species, which in this case, was a nonprimate mammal (“far”).

Other theoretically possible KGSE categories (far-near, near-N/A, far-N/A) were not observed. Indels, gaps of any type, and blanks in any nonhuman species’ alignment to human at a KGSE resulted in that species not being scored.

## 5. Conclusions

We present the first systematic analysis of PCOS-associated lncRNAs that integrates gene expression data, statistically significant GWAS SNP disease associations, FANTOM5 promoter activity, and gene-structure evolutionary conservation. Among the 23 PDEGAL lncRNAs, several exhibited disease-associated SNPs, expression in reproductive tissues, and granulosa cell transcriptional activity, highlighting them as potential functional regulators in PCOS, which also exhibit primate-specific gene structures and evidence of recent structural changes of mammalian-wide gene features in primates. These findings provide a foundation for future experimental validations, and emphasize the importance of canvassing and leveraging upon multidimensional datasets to uncover the regulatory landscape of complex diseases such as PCOS.

## Supplementary Material

**Table S1. Comprehensive dataset of candidate PDEGAL lncRNAs and associated genomic, expression, and evolutionary data**.

**Table S2. PDEGAL’s genetic, expression, and evolutionary evidence of relevance to PCOS in 4 criteria**.

**Table S3. Evolutionary classification of introns based on paired conservation analysis of splice donors and acceptors**.

**Figure S1. Phylogenetic diagram–based conservation analysis of PDE4B-AS1**.

## Author Contributions

Conceptualization, Leonard Lipovich; Data curation, Zhizhou He and Yibai Li; Formal analysis, Zhizhou He and Yibai Li; Funding acquisition, Tatiana P. Shkurat and Leonard Lipovich; Investigation, Zhizhou He and Yibai Li; Methodology, Zhizhou He, Yibai Li and Leonard Lipovich; Project administration, Li Chen and Leonard Lipovich; Supervision, Li Chen and Leonard Lipovich; Visualization, Zhizhou He and Yibai Li; Writing – original draft, Zhizhou He and Yibai Li; Writing – review & editing, Tatiana P. Shkurat, Elena V. Butenko, Ekaterina G. Derevyanchuk, Svetlana V. Lomteva, Li Chen and Leonard Lipovich.

## Data Availability Statement

Not applicable.

## Abbreviations

The following abbreviations are used in this manuscript:

PCOS: Polycystic Ovary Syndrome
lncRNA: Long Non-coding RNA
PDEGAL: PCOS differentially expressed gene associated lncRNAs
SNP: Single Nucleotide Polymorphism
GWAS: genome-wide association study
GTEx: Genotype-Tissue Expression
TSS: transcription start sites
CAGE: cap analysis of gene expression
CTSS: CAGE Transcription Start Sites
BAM: Binary Alignment Map
TPM: transcripts per million
KGSE: Key Gene Structure Elements

## References

1. Balen, A.H.; Morley, L.C.; Misso, M.; Franks, S.; Legro, R.S.; Wijeyaratne, C.N.; Stener-Victorin, E.; Fauser, B.C.; Norman, R.J.; Teede, H. The management of anovulatory infertility in women with polycystic ovary syndrome: an analysis of the evidence to support the development of global WHO guidance. Hum Reprod Update 2016, 22, 687–708, doi:10.1093/humupd/dmw025.

2. Deswal, R.; Narwal, V.; Dang, A.; Pundir, C.S. The Prevalence of Polycystic Ovary Syndrome: A Brief Systematic Review. J Hum Reprod Sci 2020, 13, 261–271, doi:10.4103/jhrs.JHRS_95_18.

3. Azziz, R. PCOS in 2015: New insights into the genetics of polycystic ovary syndrome. Nat Rev Endocrinol 2016, 12, 183, doi:10.1038/nrendo.2016.9.

4. Escobar-Morreale, H.F. Polycystic ovary syndrome: definition, aetiology, diagnosis and treatment. Nat Rev Endocrinol 2018, 14, 270–284, doi:10.1038/nrendo.2018.24.

5. Garruti, G.; Depalo, R.; Vita, M.G.; Lorusso, F.; Giampetruzzi, F.; Damato, A.B.; Giorgino, F. Adipose tissue, metabolic syndrome and polycystic ovary syndrome: from pathophysiology to treatment. Reprod Biomed Online 2009, 19, 552–563, doi:10.1016/j.rbmo.2009.05.010.

6. Randeva, H.S.; Tan, B.K.; Weickert, M.O.; Lois, K.; Nestler, J.E.; Sattar, N.; Lehnert, H. Cardiometabolic aspects of the polycystic ovary syndrome. Endocr Rev 2012, 33, 812–841, doi:10.1210/er.2012-1003.

7. Boomsma, C.M.; Eijkemans, M.J.; Hughes, E.G.; Visser, G.H.; Fauser, B.C.; Macklon, N.S. A meta-analysis of pregnancy outcomes in women with polycystic ovary syndrome. Hum Reprod Update 2006, 12, 673–683, doi:10.1093/humupd/dml036.

8. Palomba, S.; La Sala, G.B. Pregnancy complications in women with polycystic ovary syndrome: importance of diagnostic criteria or of phenotypic features? Hum Reprod 2016, 31, 223–224, doi:10.1093/humrep/dev284.

9. Qin, J.Z.; Pang, L.H.; Li, M.J.; Fan, X.J.; Huang, R.D.; Chen, H.Y. Obstetric complications in women with polycystic ovary syndrome: a systematic review and meta-analysis. Reprod Biol Endocrinol 2013, 11, 56, doi:10.1186/1477-7827-11-56.

10. Matsuda, F.; Inoue, N.; Manabe, N.; Ohkura, S. Follicular growth and atresia in mammalian ovaries: regulation by survival and death of granulosa cells. J Reprod Dev 2012, 58, 44–50, doi:10.1262/jrd.2011-012.

11. Shen, H.; Wang, Y. Activation of TGF-beta1/Smad3 signaling pathway inhibits the development of ovarian follicle in polycystic ovary syndrome by promoting apoptosis of granulosa cells. J Cell Physiol 2019, 234, 11976–11985, doi:10.1002/jcp.27854.

12. De Diego, M.V.; Gomez-Pardo, O.; Groar, J.K.; Lopez-Escobar, A.; Martin-Estal, I.; Castilla-Cortazar, I.; Rodriguez-Zambrano, M.A. Metabolic impact of current therapeutic strategies in Polycystic Ovary Syndrome: a preliminary study. Arch Gynecol Obstet 2020, 302, 1169–1179, doi:10.1007/s00404-020-05696-y.

13. Jin, P.; Xie, Y. Treatment strategies for women with polycystic ovary syndrome. Gynecol Endocrinol 2018, 34, 272–277, doi:10.1080/09513590.2017.1395841.

14. Derrien, T.; Johnson, R.; Bussotti, G.; Tanzer, A.; Djebali, S.; Tilgner, H.; Guernec, G.; Martin, D.; Merkel, A.; Knowles, D.G.; et al. The GENCODE v7 catalog of human long noncoding RNAs: analysis of their gene structure, evolution, and expression. Genome Res 2012, 22, 1775–1789, doi:10.1101/gr.132159.111.

15. Yang, D.; Wang, Y.; Zheng, Y.; Dai, F.; Liu, S.; Yuan, M.; Deng, Z.; Bao, A.; Cheng, Y. Silencing of lncRNA UCA1 inhibited the pathological progression in PCOS mice through the regulation of PI3K/AKT signaling pathway. J Ovarian Res 2021, 14, 48, doi:10.1186/s13048-021-00792-2.

16. Mudge, J.M.; Carbonell-Sala, S.; Diekhans, M.; Martinez, J.G.; Hunt, T.; Jungreis, I.; Loveland, J.E.; Arnan, C.; Barnes, I.; Bennett, R.; et al. GENCODE 2025: reference gene annotation for human and mouse. Nucleic Acids Res 2025, 53, D966–D975, doi:10.1093/nar/gkae1078.

17. Carninci, P.; Kasukawa, T.; Katayama, S.; Gough, J.; Frith, M.C.; Maeda, N.; Oyama, R.; Ravasi, T.; Lenhard, B.; Wells, C.; et al. The transcriptional landscape of the mammalian genome. Science 2005, 309, 1559–1563, doi:10.1126/science.1112014.

18. Engstrom, P.G.; Suzuki, H.; Ninomiya, N.; Akalin, A.; Sessa, L.; Lavorgna, G.; Brozzi, A.; Luzi, L.; Tan, S.L.; Yang, L.; et al. Complex Loci in human and mouse genomes. PLoS Genet 2006, 2, e47, doi:10.1371/journal.pgen.0020047.

19. Banfai, B.; Jia, H.; Khatun, J.; Wood, E.; Risk, B.; Gundling, W.E., Jr.; Kundaje, A.; Gunawardena, H.P.; Yu, Y.; Xie, L.; et al. Long noncoding RNAs are rarely translated in two human cell lines. Genome Res 2012, 22, 1646–1657, doi:10.1101/gr.134767.111.

20. Heintzman, N.D.; Stuart, R.K.; Hon, G.; Fu, Y.; Ching, C.W.; Hawkins, R.D.; Barrera, L.O.; Van Calcar, S.; Qu, C.; Ching, K.A.; et al. Distinct and predictive chromatin signatures of transcriptional promoters and enhancers in the human genome. Nat Genet 2007, 39, 311–318, doi:10.1038/ng1966.

21. Ledford, H. Circular RNAs throw genetics for a loop. Nature 2013, 494, 415, doi:10.1038/494415a.

22. Orom, U.A.; Derrien, T.; Beringer, M.; Gumireddy, K.; Gardini, A.; Bussotti, G.; Lai, F.; Zytnicki, M.; Notredame, C.; Huang, Q.; et al. Long noncoding RNAs with enhancer-like function in human cells. Cell 2010, 143, 46–58, doi:10.1016/j.cell.2010.09.001.

23. Piergentili, R.; Sechi, S.; De Paola, L.; Zaami, S.; Marinelli, E. Building a Hand-Curated ceRNET for Endometrial Cancer, Striving for Clinical as Well as Medicolegal Soundness: A Systematic Review. Noncoding RNA 2025, 11, doi:10.3390/ncrna11030034.

24. Katayama, S.; Tomaru, Y.; Kasukawa, T.; Waki, K.; Nakanishi, M.; Nakamura, M.; Nishida, H.; Yap, C.C.; Suzuki, M.; Kawai, J.; et al. Antisense transcription in the mammalian transcriptome. Science 2005, 309, 1564–1566, doi:10.1126/science.1112009.

25. Chen, X.; He, H.; Long, B.; Wei, B.; Yang, P.; Huang, X.; Wang, Q.; Lin, J.; Tang, H. Acupuncture regulates the apoptosis of ovarian granulosa cells in polycystic ovarian syndrome-related abnormal follicular development through LncMEG3-mediated inhibition of miR-21-3p. Biol Res 2023, 56, 31, doi:10.1186/s40659-023-00441-6.

26. Dykes, I.M.; Emanueli, C. Transcriptional and Post-transcriptional Gene Regulation by Long Non-coding RNA. Genomics Proteomics Bioinformatics 2017, 15, 177–186, doi:10.1016/j.gpb.2016.12.005.

27. Lin, C.Y.; Kleinbrink, E.L.; Dachet, F.; Cai, J.; Ju, D.; Goldstone, A.; Wood, E.J.; Liu, K.; Jia, H.; Goustin, A.S.; et al. Primate-specific oestrogen-responsive long non-coding RNAs regulate proliferation and viability of human breast cancer cells. Open Biol 2016, 6, doi:10.1098/rsob.150262.

28. Mittal, P.; Romero, R.; Tarca, A.L.; Gonzalez, J.; Draghici, S.; Xu, Y.; Dong, Z.; Nhan-Chang, C.L.; Chaiworapongsa, T.; Lye, S.; et al. Characterization of the myometrial transcriptome and biological pathways of spontaneous human labor at term. J Perinat Med 2010, 38, 617–643, doi:10.1515/jpm.2010.097.

29. Romero, R.; Tarca, A.L.; Chaemsaithong, P.; Miranda, J.; Chaiworapongsa, T.; Jia, H.; Hassan, S.S.; Kalita, C.A.; Cai, J.; Yeo, L.; et al. Transcriptome interrogation of human myometrium identifies differentially expressed sense-antisense pairs of protein-coding and long non-coding RNA genes in spontaneous labor at term. J Matern Fetal Neonatal Med 2014, 27, 1397–1408, doi:10.3109/14767058.2013.860963.

30. Kleinbrink, E.L.; Gomez-Lopez, N.; Ju, D.; Done, B.; Goustin, A.S.; Tarca, A.L.; Romero, R.; Lipovich, L. Gestational Age Dependence of the Maternal Circulating Long Non-Coding RNA Transcriptome During Normal Pregnancy Highlights Antisense and Pseudogene Transcripts. Front Genet 2021, 12, 760849, doi:10.3389/fgene.2021.760849.

31. Cox, M.J.; Edwards, M.C.; Rodriguez Paris, V.; Aflatounian, A.; Ledger, W.L.; Gilchrist, R.B.; Padmanabhan, V.; Handelsman, D.J.; Walters, K.A. Androgen Action in Adipose Tissue and the Brain are Key Mediators in the Development of PCOS Traits in a Mouse Model. Endocrinology 2020, 161, doi:10.1210/endocr/bqaa061.

32. Qi, X.; Zhang, B.; Zhao, Y.; Li, R.; Chang, H.M.; Pang, Y.; Qiao, J. Hyperhomocysteinemia Promotes Insulin Resistance and Adipose Tissue Inflammation in PCOS Mice Through Modulating M2 Macrophage Polarization via Estrogen Suppression. Endocrinology 2017, 158, 1181–1193, doi:10.1210/en.2017-00039.

33. Song, B.S.; Kim, J.S.; Kim, Y.H.; Sim, B.W.; Yoon, S.B.; Cha, J.J.; Choi, S.A.; Yang, H.J.; Mun, S.E.; Park, Y.H.; et al. Induction of autophagy during in vitro maturation improves the nuclear and cytoplasmic maturation of porcine oocytes. Reprod Fertil Dev 2014, 26, 974–981, doi:10.1071/RD13106.

34. Diao, F.Y.; Xu, M.; Hu, Y.; Li, J.; Xu, Z.; Lin, M.; Wang, L.; Zhou, Y.; Zhou, Z.; Liu, J.; et al. The molecular characteristics of polycystic ovary syndrome (PCOS) ovary defined by human ovary cDNA microarray. J Mol Endocrinol 2004, 33, 59–72, doi:10.1677/jme.0.0330059.

35. Jia, H.; Osak, M.; Bogu, G.K.; Stanton, L.W.; Johnson, R.; Lipovich, L. Genome-wide computational identification and manual annotation of human long noncoding RNA genes. RNA 2010, 16, 1478–1487, doi:10.1261/rna.1951310.

36. Lipovich, L.; Tarca, A.L.; Cai, J.; Jia, H.; Chugani, H.T.; Sterner, K.N.; Grossman, L.I.; Uddin, M.; Hof, P.R.; Sherwood, C.C.; et al. Developmental changes in the transcriptome of human cerebral cortex tissue: long noncoding RNA transcripts. Cereb Cortex 2014, 24, 1451–1459, doi:10.1093/cercor/bhs414.

37. Butenko, E.V.; Lipovich, L. Bioinformatic search for miRNA and lncRNA genes around genes involved in the pathogenesis of human polycystic ovary syndrome. Living and Bio-inert Systems 2024, Article 9, doi:10.18522/2308-9709-2024-49-9.

38. Severin, J.; Lizio, M.; Harshbarger, J.; Kawaji, H.; Daub, C.O.; Hayashizaki, Y.; Bertin, N.; Forrest, A.R. Interactive visualization and analysis of large-scale sequencing datasets using ZENBU. Nat Biotechnol 2014, 32, 217–219, doi:10.1038/nbt.2840.

39. Brosius, J.; Gould, S.J. On “genomenclature”: a comprehensive (and respectful) taxonomy for pseudogenes and other “junk DNA”. Proc Natl Acad Sci U S A 1992, 89, 10706–10710, doi:10.1073/pnas.89.22.10706.

40. Wood, E.J.; Chin-Inmanu, K.; Jia, H.; Lipovich, L. Sense-antisense gene pairs: sequence, transcription, and structure are not conserved between human and mouse. Front Genet 2013, 4, 183, doi:10.3389/fgene.2013.00183.

41. Kent, W.J.; Sugnet, C.W.; Furey, T.S.; Roskin, K.M.; Pringle, T.H.; Zahler, A.M.; Haussler, D. The human genome browser at UCSC. Genome Res 2002, 12, 996–1006, doi:10.1101/gr.229102.

42. Perez, G.; Barber, G.P.; Benet-Pages, A.; Casper, J.; Clawson, H.; Diekhans, M.; Fischer, C.; Gonzalez, J.N.; Hinrichs, A.S.; Lee, C.M.; et al. The UCSC Genome Browser database: 2025 update. Nucleic Acids Res 2025, 53, D1243–D1249, doi:10.1093/nar/gkae974.

43. Frankish, A.; Diekhans, M.; Ferreira, A.M.; Johnson, R.; Jungreis, I.; Loveland, J.; Mudge, J.M.; Sisu, C.; Wright, J.; Armstrong, J.; et al. GENCODE reference annotation for the human and mouse genomes. Nucleic Acids Res 2019, 47, D766–D773, doi:10.1093/nar/gky955.

44. O’Leary, N.A.; Wright, M.W.; Brister, J.R.; Ciufo, S.; Haddad, D.; McVeigh, R.; Rajput, B.; Robbertse, B.; Smith-White, B.; Ako-Adjei, D.; et al. Reference sequence (RefSeq) database at NCBI: current status, taxonomic expansion, and functional annotation. Nucleic Acids Res 2016, 44, D733–745, doi:10.1093/nar/gkv1189.

45. Pruitt, K.D.; Tatusova, T.; Maglott, D.R. NCBI Reference Sequence (RefSeq): a curated non-redundant sequence database of genomes, transcripts and proteins. Nucleic Acids Res 2005, 33, D501–504, doi:10.1093/nar/gki025.

46. Buniello, A.; MacArthur, J.A.L.; Cerezo, M.; Harris, L.W.; Hayhurst, J.; Malangone, C.; McMahon, A.; Morales, J.; Mountjoy, E.; Sollis, E.; et al. The NHGRI-EBI GWAS Catalog of published genome-wide association studies, targeted arrays and summary statistics 2019. Nucleic Acids Res 2019, 47, D1005–D1012, doi:10.1093/nar/gky1120.

47. ENCODE Consortium co-author list. The GTEx Consortium atlas of genetic regulatory effects across human tissues. Science 2020, 369, 1318–1330, doi:10.1126/science.aaz1776.

48. Blanchette, M.; Kent, W.J.; Riemer, C.; Elnitski, L.; Smit, A.F.; Roskin, K.M.; Baertsch, R.; Rosenbloom, K.; Clawson, H.; Green, E.D.; et al. Aligning multiple genomic sequences with the threaded blockset aligner. Genome Res 2004, 14, 708–715, doi:10.1101/gr.1933104.

49. FANTOM Consortium co-author list including Lipovich L. A promoter-level mammalian expression atlas. Nature 2014, 507, 462–470, doi:10.1038/nature13182.

50. Hon, C.C.; Ramilowski, J.A.; Harshbarger, J.; Bertin, N.; Rackham, O.J.; Gough, J.; Denisenko, E.; Schmeier, S.; Poulsen, T.M.; Severin, J.; et al. An atlas of human long non-coding RNAs with accurate 5’ ends. Nature 2017, 543, 199–204, doi:10.1038/nature21374.

