## Supplementary material for "Integrative Identification and Characterization of PCOS-Associated lncRNAs From the Interface of Genetic Association, Transcriptomics, and Gene Structure Evolution": Figure S1.docx

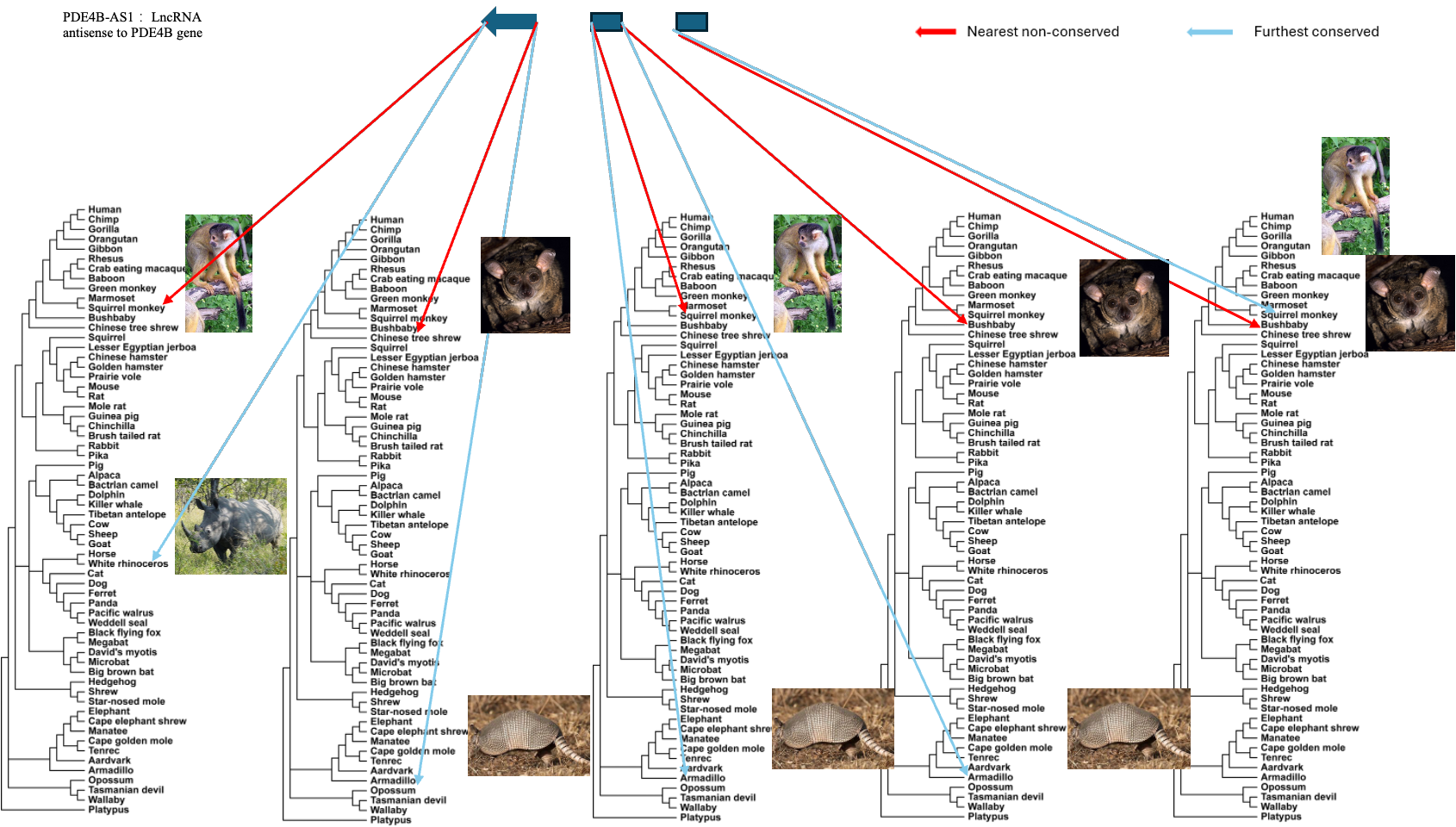

**Figure S1. Phylogenetic diagram–based conservation analysis of PDE4B-AS1.** Two selected key gene structure elements (KGSEs) of PDE4B-AS1 (intron 1 splice donor GT- and intron 1 splice acceptor -AG) were analyzed for evolutionary conservation across mammals in the 99-species MultiZ alignment. The phylogenetic tree topology and representative species images were adapted from the UCSC Genome Browser Conservation track (99-species MultiZ alignment). The final trees were rendered and arranged using iTOL (Interactive Tree Of Life, https://itol.embl.de/). Red arrows indicate the nearest species (to human) where the KGSE is non-conserved, and blue arrows indicate the furthest species (from human) where the KGSE is conserved.
